# AMP-activated-protein kinase (AMPK) is an essential sensor and metabolic regulator of retinal neurons and their integrated metabolism with RPE

**DOI:** 10.1101/2020.05.22.109165

**Authors:** Lei Xu, Emily E. Brown, Casey J. Keuthan, Himani Gubbi, Elodie-Kim Grellier, Jerome Roger, Anand Swaroop, Jianhai Du, John D. Ash

**Affiliations:** Department of Ophthalmology, College of Medicine, University of Florida, Gainesville, FL 32610 USA; Paris-Saclay Institute of Neuroscience, CERTO-Retina France, CNRS, Univeriste Paris-Saclay, Orsay, France; Neurobiology Neurodegeneration & Repair Laboratory, National Eye Institute, NIH, Bethesda, Maryland; Department of Ophthalmology and Visual Sciences, Department of Biochemistry, School of Medicine, West Virginia University, Morgantown, WV 26506

## Abstract

The retina has one of the highest energy demands in the human body, and proper regulation of metabolism among the cell types of the retina is required for functional vision. Recent studies have reported that retinal metabolism is disrupted during retinal degeneration and aging, while therapies designed to enhance metabolism can slow or prevent degeneration. Here, we show that multiple metabolic pathways, including those regulated by a key ATP sensor and regulator of metabolism, AMP-activated-kinase (AMPK), were dysregulated in a model of inherited retinal degeneration. In order to assess the direct role of AMPK in regulating retinal metabolism, we used mice with retina-specific knockout of AMPK activity. Conditional loss of AMPK resulted in impaired visual function at an early age, with slow photoreceptor loss observed in older mice. Moreover, we found that loss of AMPK resulted in decreased metabolic flux from glucose, decreased mitochondrial DNA copy number, decreased mitochondrial-related gene expression, and alterations in mitochondrial morphology in the photoreceptors, all of which preceded degeneration. Surprisingly, metabolic changes from the loss of AMPK in retinal neurons also resulted in secondary degeneration of retinal pigment epithelial (RPE) cells. Together, these data show that AMPK signaling is important for maintaining metabolic homeostasis of the retina and support the hypothesis that photoreceptors and RPE have a shared metabolic ecosystem that is controlled, in part, by AMPK.

---

The central nervous system, including the retina, is often cited as having one of the highest energy demands in the body, with pathways that require utilization of ATP, glucose, lactate, fatty acids, glutathione (GSH), and NADPH for proper neuronal function and protection against retinal degeneration ^1–4^. As much as half of the metabolites and metabolic activity in the retina are located in the light-sensitive photoreceptors ^5,6^. ATP is heavily consumed by NaK-ATPases in photoreceptors to maintain the ion gradient within the cells, and a considerable amount of energy is dedicated to supporting phototransduction in response to light ^1,2^. While disruption of metabolism has been associated with retinal degeneration, manipulations that restore energy homeostasis by enhancing glycolysis or mitochondrial activity can preserve retinal function and promote photoreceptor survival ^7–12^.

However, little is known about how metabolic activity is regulated in the retina. The 5’ adenosine monophosphate-activated protein kinase (AMPK) pathway is a logical control mechanism for high metabolic activity in cells. AMPK is an evolutionarily conserved serine/threonine kinase that functions as a heterotrimeric protein comprised of a single catalytic α-subunit (encoded by either *Prkaa1* or *Prkaa2* in mice), a regulatory β-regulatory subunit (encoded by either *Prkab1* or *Prkab2* in mice), and the AMP-binding *γ*-subunit (encoded by *Prkag1, Prkag2*, or *Prkag3* in mice)^13^. Expression levels of each of the AMPK subunit isoforms has previously been characterized in both human and mouse retinas^9^. AMPK is activated by increased AMP:ATP ratios within cells, which only occurs with ATP depletion. Thus, AMPK serves as a cellular energy sensor in response to nutrient deprivation, low energy states, fasting, hypoxia, as well as other stress conditions ^14,15^. Once activated, AMPK restores the energy balance by increasing glycolysis and mitochondrial biogenesis, while also reducing energy use by decreasing protein and fatty acid synthesis ^14,16–28^. This combined effect on energy expenditure and energy production signifies major metabolic reprogramming within the cells. We have previously demonstrated that stimulation of AMPK with metformin protected both photoreceptors and retinal pigment epithelial (RPE) cells in multiple mouse models of retinal degeneration, suggesting that AMPK can restore metabolic homeostasis in these cells^9^.

Here we demonstrate that AMPK functions as a key regulator of metabolism in the retina. In this study, we discovered that metabolic gene expression, including many components of the AMPK pathway, was significantly disrupted in photoreceptors prior to cell death caused by a mutation that leads to progressive, inherited retinal degeneration. We show that activation of AMPK with metformin enhanced metabolism in the healthy neuroretina, while loss of AMPK activity reduced the retina’s glycolytic and mitochondrial metabolic state. This metabolic shift due to the loss of AMPK coincided with an early decline in neuroretinal function, followed by an age-related acceleration of photoreceptor death. Interestingly, we found that ablation of AMPK in the neuroretina also resulted in an RPE degenerative phenotype in these mice, supporting the hypothesis that AMPK plays a central role in the coordinated metabolism between the photoreceptors and the RPE.

## Results

### AMPK signaling is dysregulated in early retinal disease

We have previously reported that stimulation of AMPK with metformin protects mouse photoreceptors from the inherited *Pde6b*^*rd10*^ (*rd10*) mutation and from light damage. Metformin induced-protection required AMPK expression in the retina, suggesting that AMPK plays a crucial role in this process ^15^. These results suggest that retinal degeneration is associated with altered metabolic regulation, which could be corrected by activation of AMPK. To assess dysregulation of metabolism in *rd10* mice, we analyzed RNA-sequencing data from wild-type (WT) and *rd10* rods of mice at age P19. Mice homozygous for this mutation exhibit progressive rod degeneration starting around P19 (Figure 1a)^29–31^. Therefore, this age represents a time where the photoreceptor-specific transcriptional responses prior to rapid degeneration can be assessed.

**Figure 1.**
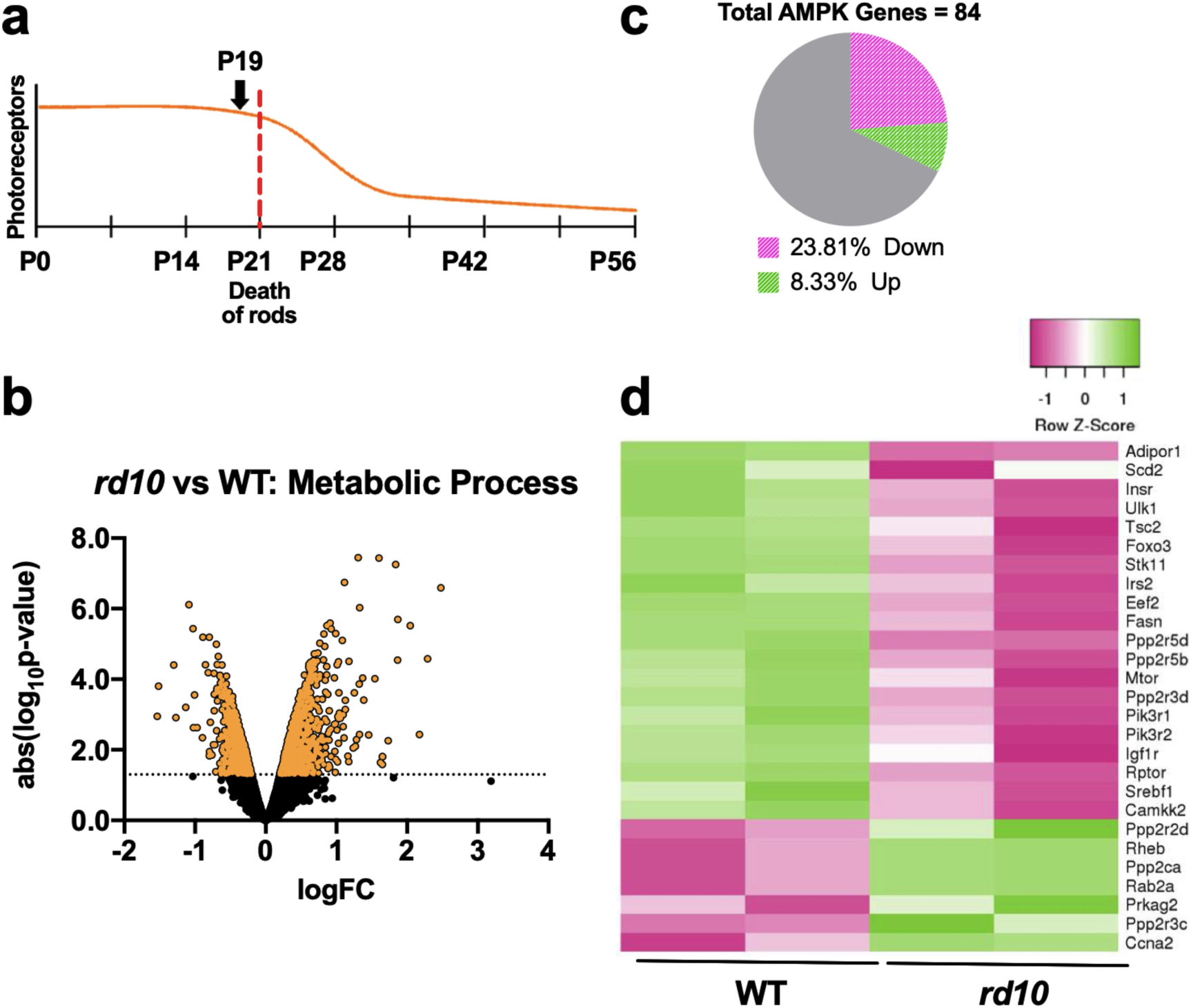
AMPK signaling is dysregulated in early inherited retinal degeneration. a) Timeline of photoreceptor degeneration in the *rd10* mouse model. Rod degeneration begins at age P21 and by age P28 few rod photoreceptors remain. RNA-seq data of *rd10* and WT rods was analyzed at age P19, prior to cell death caused by the *Pde6b*^*rd10*^ mutation. b) Volcano plot showing fold change and significance of metabolism-related gene expression detected in *rd10* rods compared to age-matched controls. Orange dots indicate a differentially expressed gene (p < 0.05). Significance threshold is also indicated by the dotted line. c) AMPK signaling genes differentially expressed in *rd10* rods. Of the 84 AMPK genes detected, over 28% were dysregulated in the mutant photoreceptors. d) Clustered heatmap of differentially expressed AMPK signaling genes in *rd10* versus WT rods. Pink indicates downregulated or lower expression, green indicates upregulated or higher expression.

To determine how metabolism was altered early in degeneration. we first mined the Gene Ontology (GO) resource to construct a comprehensive panel of genes involved specifically in metabolism ^32,33^. “Metabolic process” (GO:0008152) is a broad or high-level GO class, defined as the reactions and pathways by which living organisms transform chemical substances. This large biological program is comprised of 10,433 genes encompassing 3,331 unique GO nodes, that include a variety of anabolic and catabolic processes for the transformation of small molecules, as well as pathways for DNA repair, protein synthesis, and protein degradation. Of the genes included in this reference list, 5,857 of the metabolism genes had detectable expression levels in the rod dataset (Supplemental Table 1). Differential expression analysis of these genes revealed that 1,585 or 27.1% were dysregulated in *rd10* rods at this early time point (Figure 1b and Supplemental Table 1). These findings suggest that there is metabolic reprogramming occurring in photoreceptors early in the progression of retinal disease.

To further identify metabolic pathways that are disrupted in degeneration, we analyzed changes in gene expression using DAVID^34,35^ to assign dysregulated genes to their associated KEGG pathways. This database represents a collection of biochemical pathway maps that allow the visualization of interactions and examine genes within specific metabolic networks. Interestingly, AMPK signaling (mmu04152) was significantly enriched in the *rd10* dataset. The AMPK pathway is associated with 126 genes, both upstream and downstream, of the energy sensor (Supplemental Table 2). Analysis of this subset of genes revealed that 84 AMPK pathway genes were expressed in rods and that 27 were significantly changed in the diseased photoreceptors (Figure 1c and Supplemental Table 2). *Prkag2*, encoding one of the gamma subunits for AMPK, was upregulated, while many of the upstream activators of AMPK were decreased in *rd10* rods, including genes encoding liver kinase B1 (*Stk11*) and CAM-kinase kinase (*Camkk2*) (Figure 1d). Other components of insulin receptor signaling (e.g. *Insr*, *Igf1r*) that lead to AMPK activation were similarly decreased (Figure 1d). Moreover, *rd10* rods had significant decreases in expression of downstream AMPK targets, including master regulators like *Srebf1* and *Foxo3* (Figure 1d).

### Metformin treatment induces metabolic changes in the retina

Our previous data demonstrated that treatment with metformin protected mouse photoreceptors from degeneration, likely due to metabolic reprogramming of the retinal cells. To determine how metformin alters retinal metabolism, we measured steady state levels of 183 metabolite intermediates including ATP, AMP, NAD+, NADH, hypoxanthine, xanthine, creatinine, phosphocreatine, cGMP, GSH, glutathione disulfide (GSSG), adenosine, the tricarboxylic acid (TCA) cycle intermediates (e.g. a-ketoglutarate, malate, and succinate), amino acids involved in energy production (e.g. proline, serine, alanine, glutamate, glutamine, and aspartic acid), and glycolysis intermediates (e.g. glucose, lactate, glucose-6-phosphate, and glucose-1-phosphate) (Supplemental Table 3). By comparing metformin and PBS-treated retinas, we found that metformin treatment resulted in increased levels of metabolites such as ATP, phosphocreatine, GSH, GSSG, proline, alanine, and lactate (Figure 2). These changes indicate that metformin induces major changes in metabolic pathways, including glycolysis, mitochondrial metabolism, glutamate metabolism, NAD metabolism, and glutathione metabolism, which are essential for normal photoreceptor physiology.

**Figure 2.**
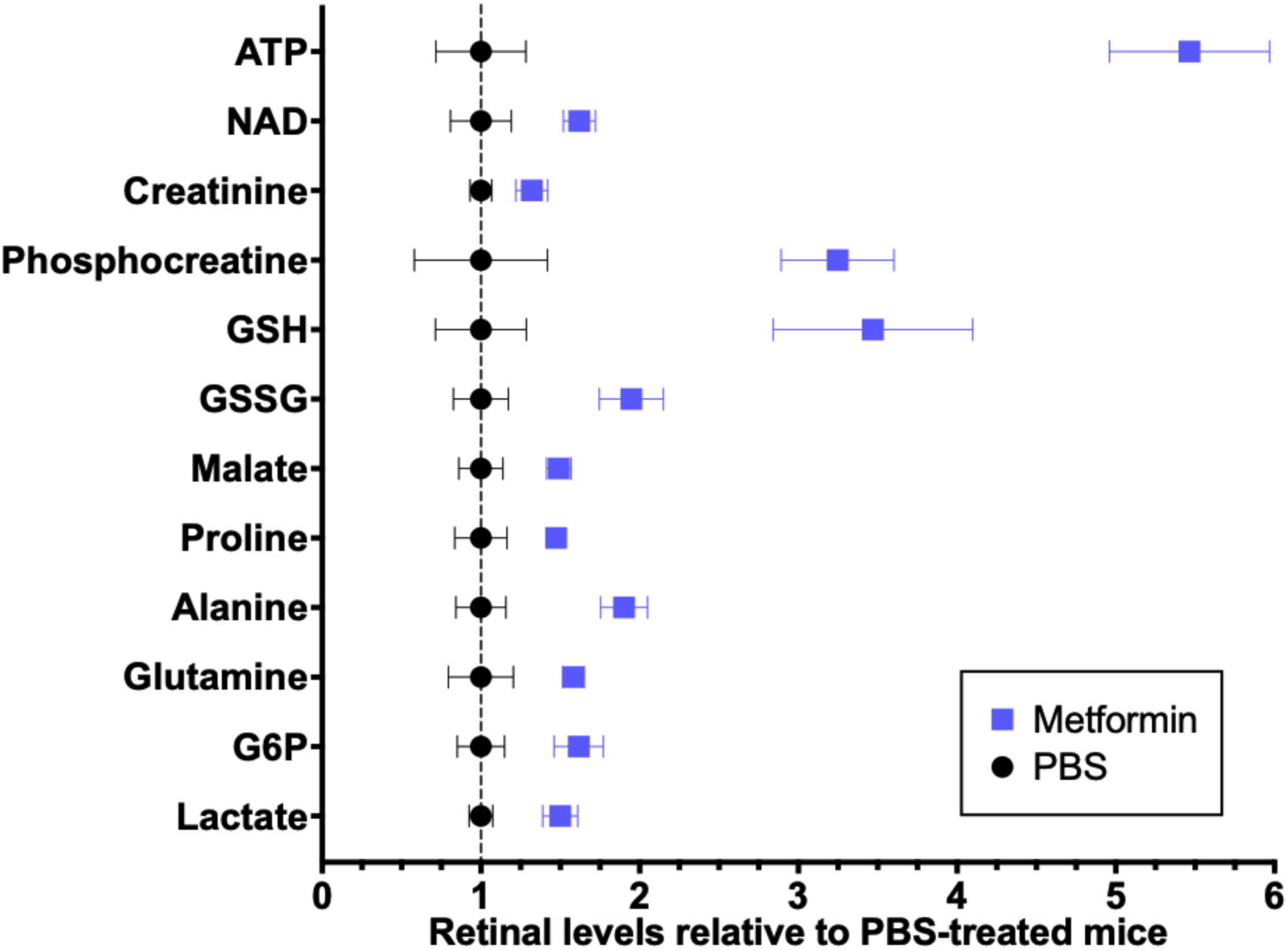
Metformin modulates retinal metabolic homeostasis. 183 metabolites were measured in metformin-treated retinas and were normalized to PBS-treated control samples (Supplemental Table 3). All metabolites with statistically significant changes are shown. Metabolites involved in energy production (ATP, NAD, creatinine, phosphocreatine), glutathione metabolism (GSH, GSSH), the TCA cycle (malate), amino acids (proline, alanine, glutamine), and glycolysis (G6P, lactate) were increased with metformin treatment. Multiple t-tests, p < 0.05, n = 3 per group.

### AMPK activity is essential for retinal morphology and function

To study the direct role of AMPK in the retina, we used a *Chx10-Cre*/loxP system to conditionally delete one (α1 KO and α2 KO) or both (also known as neuroretina double KO or nDKO) of the protein’s catalytic subunits in retinal neurons and Müller glia cells. We observed approximately a 75-80% reduction in AMPK α1 and α2 mRNA levels (encoded by *Prkaa1* and *Prkaa2*, respectively) in nDKO retinas compared to WT controls, as confirmed by qRT-PCR (Supplemental Figure 1). We initially assessed the loss of function effect of a single AMPK α subunit on retinal structure. Single AMPK α1 and α2 KO retinas appeared unremarkable as compared to age-matched WT mice up to 12 months old (Supplemental Figure 2a, c, d). However, complete loss of AMPK catalytic activity through deletion of both subunits led to more drastic effects on retinal morphology. We observed a marked reduction in total retina and inner nuclear layer (INL) thickness, as well as an obvious collapse of the photoreceptor inner segment (IS) layer and a significant loss of the outer nuclear layer (ONL) photoreceptor nuclei in 12-month-old nDKO mice (Figure 3a-d, Supplemental Figure 3). This thinning of the ONL was apparent in AMPK nDKO mice starting around 9 months of age, as confirmed by *in vivo* optical coherence tomography (OCT) measurements of retinal thickness over time (Figure 3e).

**Figure 3.**
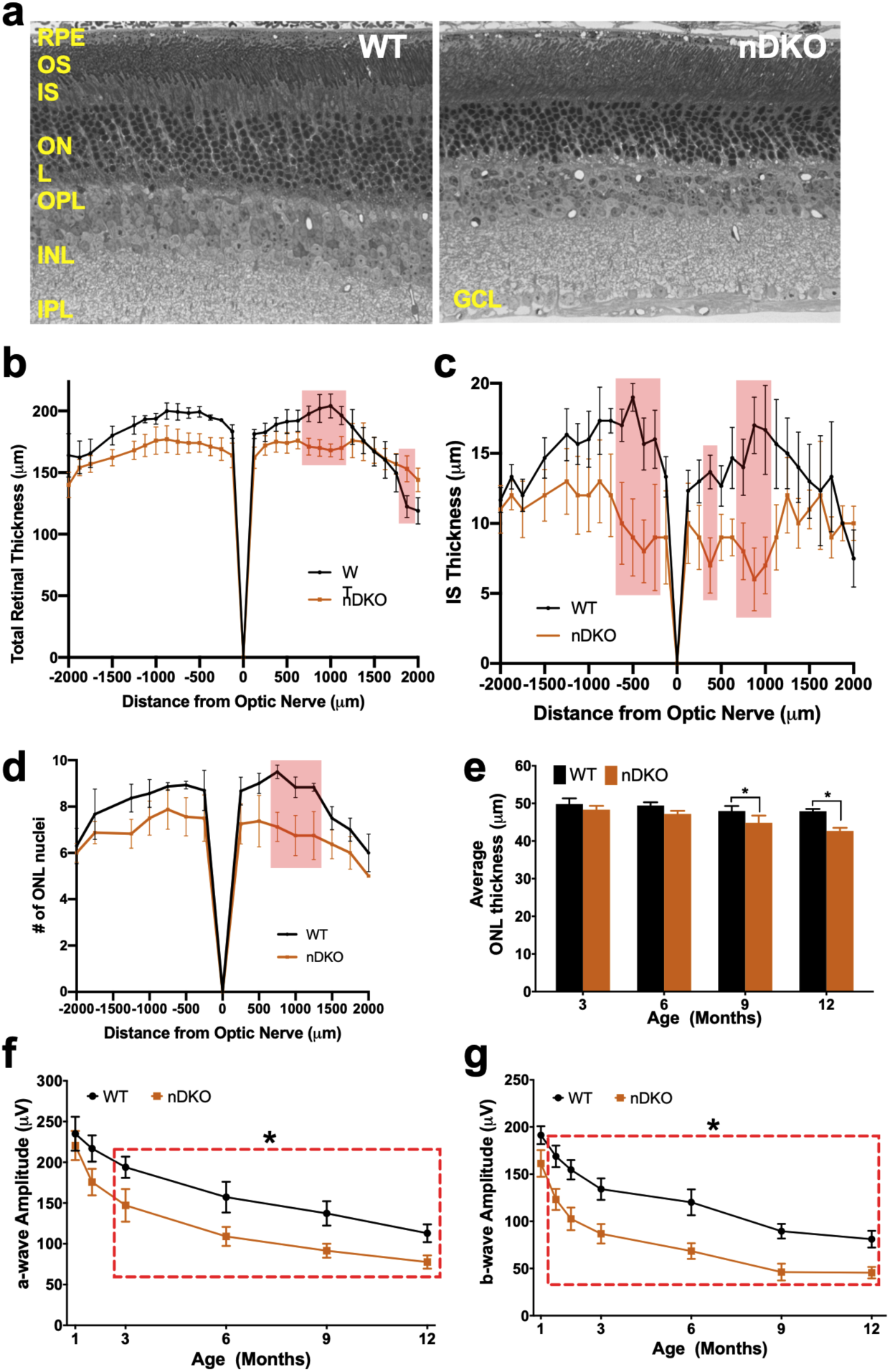
Deletion of AMPK in the neuroretina results in age-associated reduced retinal function and thinning of retinal layers. a) Histological sections of WT (left) and AMPK nDKO (right) retinas at 12 months of age. b-d) Quantification of the total retinal thickness (b), inner segment (IS) thickness (c), and number of nuclei in columns in the outer nuclear layer (ONL) (d) from 12-month-old histology sections at fixed intervals from the optic nerve. Red shaded boxes represent statistically significant values. Two-way ANOVA, p ≤ 0.05, n = 3 WT and 4 nDKO. e) Quantification of outer nuclear layer (ONL) thicknesses from OCT images with age. Two-way ANOVA, p ≤ 0.0190, n ≥ 3 per group. f) Scotopic ERG a-wave with aging. Red box indicates statistically significant values. Two-way ANOVA, p ≤ 0.0044, n≥3 per group. g) Photopic ERG b-wave with aging in AMPK nDKO or WT mice. Red box indicates statistically significant values. Two-way ANOVA, p < 0.0001, n ≥ 5 per group.

To determine the effect of AMPK on retinal function, we performed electroretinography (ERG) on nDKO and WT mice to independently assess the function of rod photoreceptors (scotopic a-wave), cone photoreceptors (photopic b-wave), and the inner retina (scotopic b-wave) over time. While deletion of the α2 subunit led to a decrease in cone function only, deletion of both AMPK α subunits resulted in a significant reduction in both rod and cone function at an early age (Supplemental Figure 2b, Figure 3f-g). The decline in cone function in nDKO mice was observed as early as 6 weeks old, followed by the loss of rod function by 3 months old (Figure 3f-g). Results with the AMPK nDKO mice suggest that AMPK activity is essential for normal retinal morphology and function. Due to the more pronounced phenotype, we decided to focus on the AMPK nDKO mice for the remainder of our study.

### Loss of AMPK significantly alters mitochondrial structure and gene expression

Given that enhanced AMPK activity leads to increased mitochondrial DNA copy number and mitochondrial-related gene expression^9^, we sought to assess changes in mitochondrial physiology in AMPK nDKO retinas. Transmission electron microscopy (TEM) images of 1-month-old nDKO mice appeared normal (Supplemental Figure 4a), which was in agreement with our findings that these mice have normal retinal morphology and function at an early age. However, disruptions in mitochondrial ultrastructure in older mice were profound. By 4 months of age, the mitochondria of AMPK nDKO retinas were highly fragmented or swollen, with a noticeable loss of cristae (Figure 4a). These findings indicate that retinal AMPK activity is important for preserving mitochondrial integrity with age.

**Figure 4.**
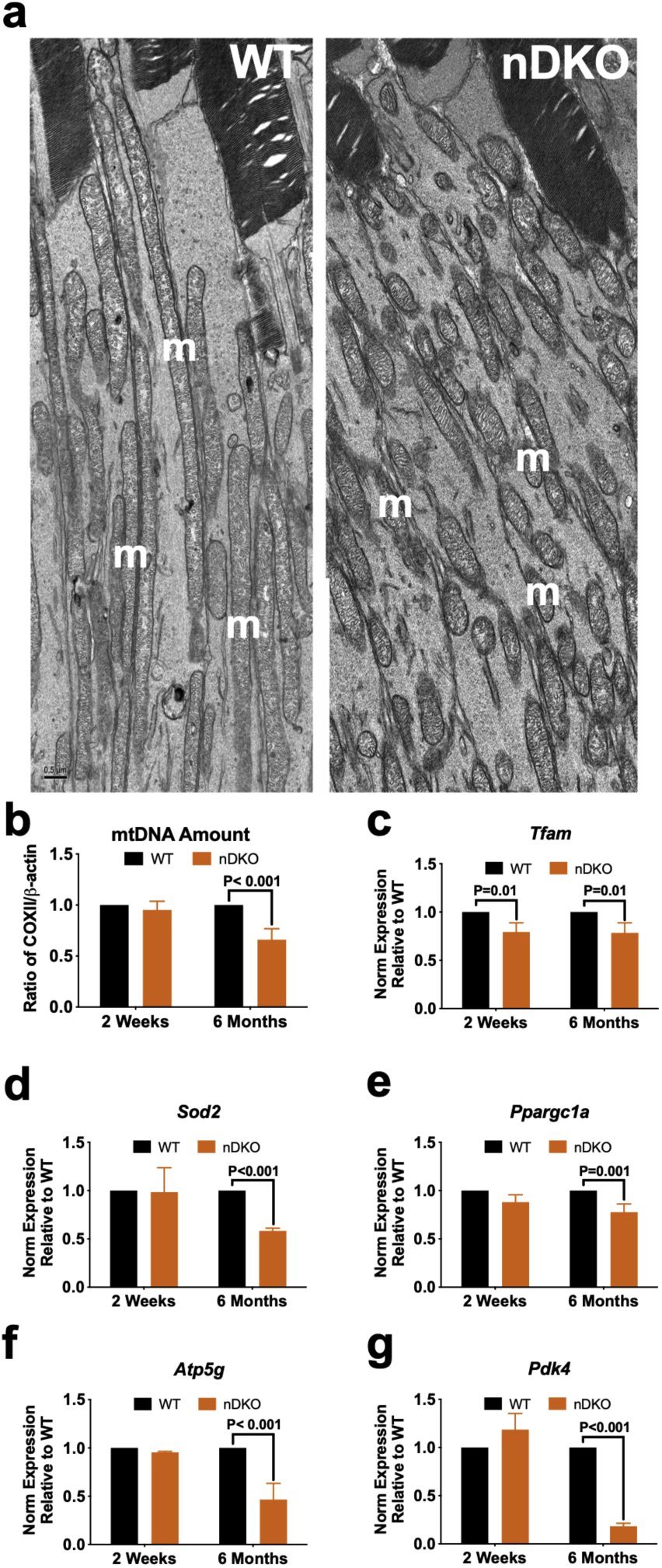
Disrupted mitochondrial morphology and gene expression in the neuroretina with deletion of AMPK. a) Representative EM images from retinal cross sections of 4 month old WT (left) and AMPK nDKO (right) mice showing the photoreceptor inner segments (IS). m, mitochondria. Scale bar indicates 0.5 µm. b-f) Gene expression analysis in the neuroretina of WT and nDKO mice at 2 weeks and 6 months of age. b) *Tfam*, a mitochondrial transcription factor. c) *Sod2*, an antioxidant enzyme located in the mitochondrial matrix. d) *Ppargac1a* (also known as PGC-1α), a master regulator of mitochondrial biogenesis. e) *Atp5g*, a component of the ATP synthase of the electron transport chain. f) *Pdk4*, an enzyme that inhibits pyruvate dehydrogenase to downregulate glucose utilization. g) Mitochondrial DNA copy number in the neuroretina at 2 weeks and 6 months of age. *CoxII* (mtDNA-encoded) divided by *Actb* (β-actin, nuclear DNA-encoded). Multiple t-tests, p-values indicated in each figure, n = 3 per group.

The mitochondrial fragmentation observed by TEM suggests that nDKO retinas may have reduced mitochondrial mass. Further analysis revealed that 6-month-old AMPK nDKO mice also had significantly less mitochondrial DNA relative to nuclear DNA content within the neuroretina as measured by qRT-PCR (Figure 4g). To further assess changes in mitochondria, we measured mRNA expression of genes involved in regulation of mitochondrial biogenesis and oxidative defense, including transcription factor A, mitochondrial (*Tfam*), peroxisome proliferator-activated receptor gamma coactivator 1-alpha (*Ppargc1a*, also known as PGC-1α), superoxide dismutase 2 (*Sod2*), membrane subunit c of the mitochondrial ATP synthase (*Atp5g*), and pyruvate dehydrogenase kinase 4 (*Pdk4*). With the exception of a slight reduction in *Tfam* expression (Figure 4b), no other genes had measurable changes in expression at 2 weeks of age (Figure 4c-f). However, by 6 months of age, there was a significant decrease in expression of *Tfam, Sod2, Ppargc1a*, *Atp5g*, and *Pdk4* in nDKO retinas as compared to WT control retinas (Figure b-e).

It is important to note that all these changes in the mitochondria occurred at an age that is prior to the loss of cells but correlated temporally with the early loss of function. Since we found reduced expression of *Sod2* in nDKO retinas, we hypothesized that AMPK-deficient retinas exhibited increased oxidative stress, which could contribute to the later-onset photoreceptor cell death in the AMPK nDKO mice. To assess this, we measured mRNA levels of thioredoxin (*Txn*), heme oxygenase 1 (*Hmox1*, also known as HO-1), and glutathione peroxidase 1 (*Gpx1*), but found no differences in expression between AMPK nDKO and WT mice (Supplemental Figure 4b-d). Together, these data suggest that retinal cells lacking AMPK activity may have insufficient energy production but are likely not in a state of elevated oxidative stress.

### Reduced retinal metabolic flux with loss of AMPK in the neuroretina

[U-^13^C] glucose is a glucose isotopomer in which all six carbons are labeled with heavy ^13^C. This allows examination of metabolic flux from glucose for metabolic processes such as glycolysis, the TCA cycle, amino acid metabolism, and glutamine metabolism. Since AMPK is a key regulator of glycolysis and mitochondrial metabolism, it is possible that a loss of AMPK activity could reduce metabolic flux in the retina through one or both of these pathways. We detected changes of 23 ^13^C-labeled metabolites between AMPK nKO and WT retinas, which were involved in glycolysis, the TCA cycle, and amino acid metabolism in 4-month-old mice (Supplemental Table 4). AMPK nDKO retinas had significant decreases in multiple glycolytic (e.g. phosphoenyl-pyruvate, pyruvate, and lactate) and TCA cycle metabolites (e.g. citrate, α-ketoglutarate, succinate, fumarate, and malate), as well as reduced labeling of aspartate, glutamate, and oxoproline. These significant changes in retinal metabolites suggest that proper regulation of metabolism is controlled through AMPK activity within retinal neurons.

Studies in other tissues have shown that AMPK can alter glucose levels through regulation of glucose transporters ^16,17^. Therefore, we hypothesized that the reduction of [U-13C] glucose labeling with AMPK loss of function may be due to altered expression levels of glucose transporter 1 (*Glut1*), which encodes for the major glucose transporter in the retina. To assess this possibility, we compared *Glut1* gene expression levels in nDKO mice with WT controls. We found normal *Glut1* expression in nDKO mice at 2 weeks of age, suggesting that *Glut1* levels are not affected during AMPK nDKO development (Figure 5b). However, by 6 months of age, *Glut1* expression in nDKO mice was significantly reduced (Figure 5b). This finding indicates that less glucose may be transported to the photoreceptors over time, which could impact glycolytic flux.

**Figure 5.**
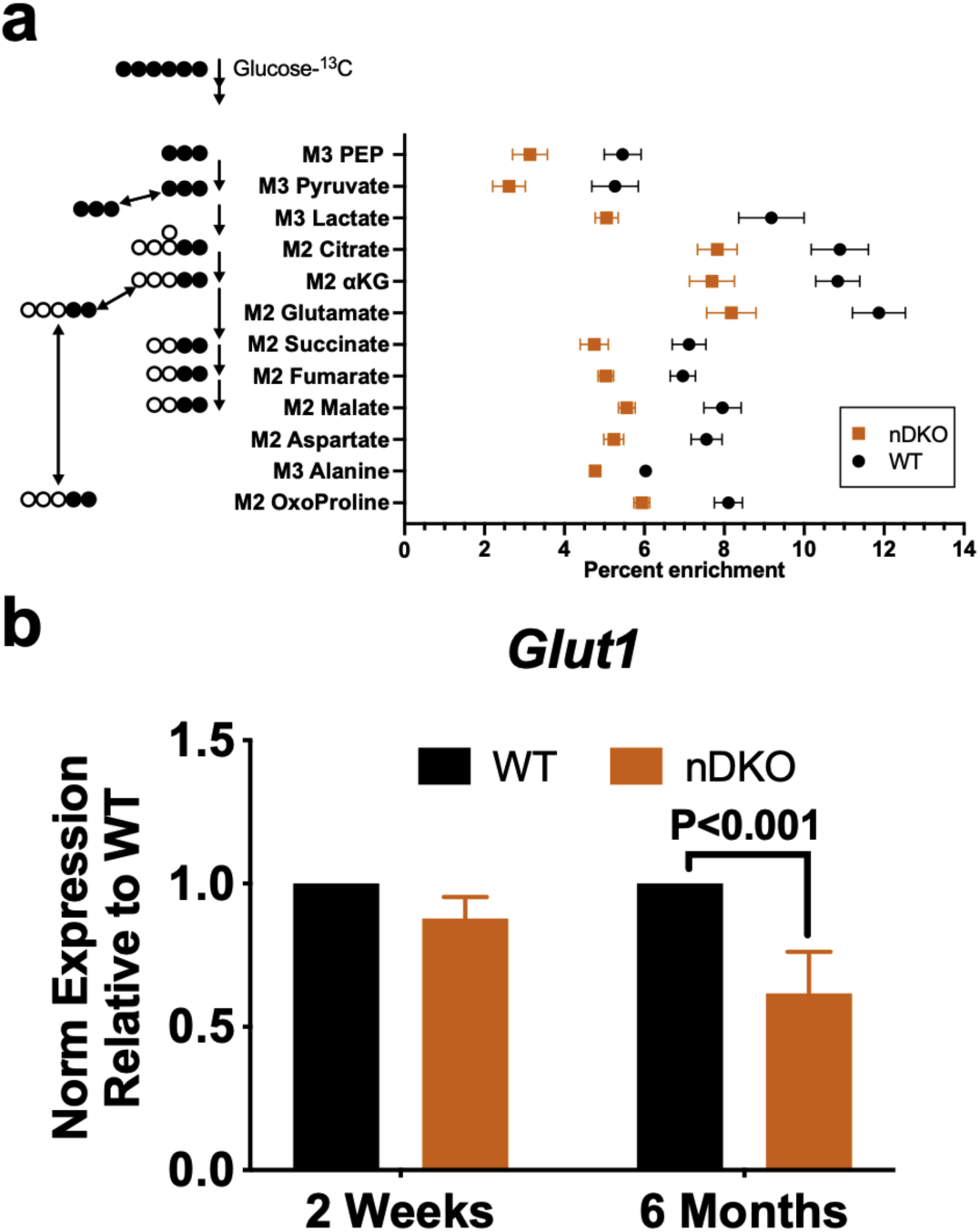
Reduced metabolic flux with deletion of AMPK in the neuroretina. a) [U-^13^C] glucose isotopologue enrichment was determined by GC-MS. Percent enrichment of metabolites with ^13^C incorporation directly from glucose in the first cycle that were statistically different are included in the graph. All isotopologues are shown in Supplemental Table 4. The label M3 or M2 in the y-axis refers to the number of labeled carbons, which are shown as closed circles in the drawing next to the y-axis labels. Multiple t-tests, n=5 per group. b) mRNA expression of glucose transporter 1 (*Glut1*) in the neuroretina in nDKO mice relative to the levels in the WT at 2 weeks and 6 months of age. Unpaired t-test, p=0.0004, n=3 per group.

### Secondary RPE abnormalities with deletion of AMPK in the neuroretina

It should be noted that our *Chx10-Cre*/loxP deletion strategy does not affect AMPK activity in astrocytes, retinal vasculature, or RPE cells. Since the RPE cells provide nourishment to the photoreceptors as part of an integrated retinal metabolism^36^, we wanted to investigate whether deletion of AMPK in the neuroretina leads to any secondary effects on the RPE. To examine RPE morphology in AMPK nDKO mice, we stained RPE flat mounts from 9-month-old mice with an antibody for zonula occludens 1 (ZO-1), a protein involved in forming RPE tight junctions. We observed a significant loss of ZO-1 staining in the nDKO mice as compared to WT controls (Figure 6a-b), suggesting that RPE architecture is affected with loss of AMPK activity in the neuroretina. We utilized TEM to further assess changes in RPE ultrastructure. Consistent with our earlier findings in the neuroretina, we did not observe any alterations in RPE ultrastructure in 1-month-old AMPK nDKO mice (Supplemental Figure 4). However, 4-month-old AMPK nDKO mice had significant RPE abnormalities, including the presence of vacuoles lacking electron density, undigested photoreceptor outer segments, lipid droplets, thickening of the RPE layer, and disorganized interdigitation of the RPE apical microvilli and photoreceptor outer segments (Figure 6c-e). These findings correlate with the reduction in retinal function observed in nDKO mice at this age. Therefore, we also measured c-wave ERG as a measure of RPE function^37^ in 12-month-old mice and found a significant reduction in the c-wave amplitude in nDKO mice as compared to that of WT controls (Figure 6f).

**Figure 6.**
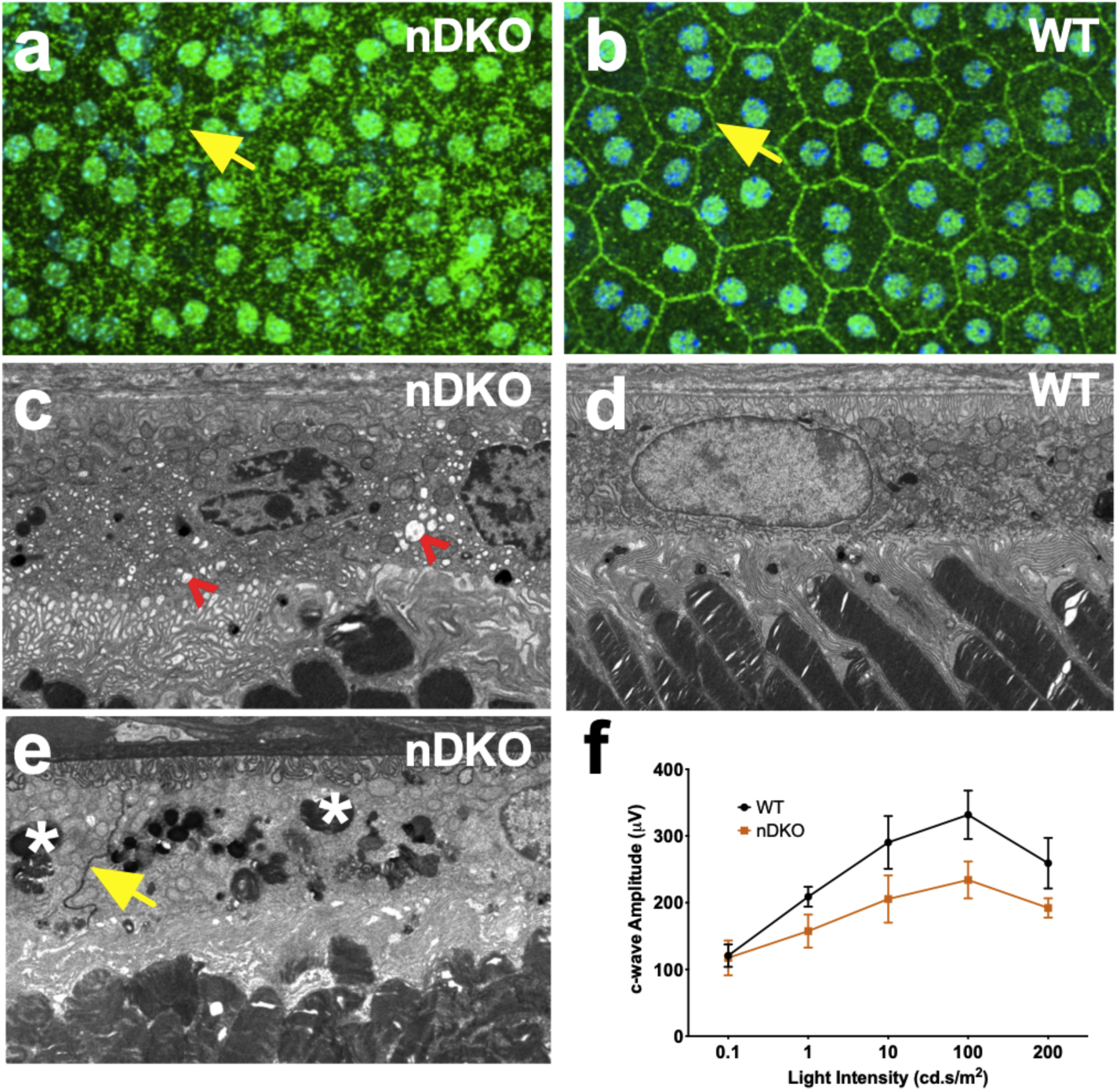
Secondary alterations in the RPE with loss of AMPK in the neuroretina. a-b) ZO-1 staining (green) and DAPI (blue) on RPE flat mounts from 9-month-old AMPK nDKO (a) and WT (b) mice. Yellow arrows denote representative tight junction cell boundaries. c-e) Representative EM images of 4-month-old AMPK nDKO (c and e) and WT (d) mice. In c, red arrows denote vacuole-like structures that lack electron density. In e, * indicate electron dense lipid, droplet-like structures and yellow arrow indicates tight junction boundary. f) RPE function as measured by c-wave ERG from 12-month-old WT and nDKO mice. Two-way ANOVA, p > 0.09, n = 6 WT and 7 nDKO.

We additionally investigated whether AMPK activity within RPE cells provided cell-autonomous regulation of metabolism. Using an RPE-specific *VMD2-Cre*, we conditionally deleted both the AMPK α1 and α2 subunits in the RPE (rDKO) to assess retinal function and morphology with age. Despite confirmation that there was approximately a 40-50% reduction in AMPK α subunit gene expression, we did not find any significant differences in retinal function or morphology with the sole loss of RPE AMPK activity (Supplemental Figure 5). Collectively, these data show that while AMPK is not necessary in the RPE under normal neuroretinal conditions, there are important, non-cell autonomous effects with the loss of AMPK exclusively in the retinal neurons.

## Discussion

We have found that AMPK plays an essential role in the regulation of metabolism in the retina and maintains the function and vitality of both the retinal neurons and the RPE. Loss of AMPK in the neuroretina led to a decline in retinal function as early as 6 weeks of age, yet cell death (loss of ONL thickness) was not observed until 9 months of age. Substantial changes in mitochondrial structure and mitochondrial-related gene expression in AMPK nDKO retinas also preceded any photoreceptor loss. Similarly, we detected widespread dysregulation of metabolism genes prior to degeneration in the *rd10* mouse model of inherited retinal disease. This was in agreement with recent metabolomic studies reporting that mice with the *Pde6b*^*rd10*^ mutation have severe disruptions in retinal metabolism ^38^. Interestingly, we found an enrichment of genes related to the AMPK pathway that were affected in the *rd10* expression dataset. Taken together, these data suggest that there are early metabolic alterations in the retina that impact cell survival, which are likely mediated by AMPK.

Activation of AMPK in the retina with metformin treatment enhanced metabolism, increasing the amount of substrates that are required for glycolysis, mitochondrial metabolism, and other key metabolic pathways. In contrast, we observed a reduction in metabolic flux through glycolytic metabolites, citric acid cycle metabolites, and metabolites involved in glutamine metabolism in AMPK nDKO retinas. Under normal conditions, 98% of glutamate in the retina is protected from catabolism by the α-ketoglutarate/glutamate circuit, which ensures that glutamate is available for neurotransmission and synthesis of glutathione, amino acids, and proteins^5^. We hypothesize that the reduction in glutamine and GABA could be due to supplementation of these metabolites into the citric acid cycle to compensate for a reduction in flux through glycolytic and citric acid cycle metabolites. Previous studies have shown that deletion of a kinase upstream from AMPK, liver kinase B1 (LKB1), or deletion of AMPK itself, in the mouse retina leads to remodeling of photoreceptor glutamatergic synapses^39^. Our data suggest that the synaptic phenotype observed previously with loss of AMPK or LKB1 in the retina may be the result of increased glutamate utilization for oxidative phosphorylation, which drives glutamate insufficiency for synaptic transmission.

Metabolic dysregulation has been linked to disease pathogenesis in both inherited retinal degenerations and age-related macular degeneration (AMD). Deletion of phosphofructokinase (PFK) in the retina accelerated retinal degeneration in a *Pde6b* mutant model of retinitis pigmentosa (RP)^8^. Other studies have shown that genes related to cellular metabolism are enriched in the cones of mouse models for inherited retinal degeneration^40^. Studies have also shown that glucose is essential for proper photoreceptor function, as deletion of GLUT1 in the RPE did not affect RPE function but led to photoreceptor cell death^41^. In addition to glycolysis, mitochondrial function is also essential for proper photoreceptor function, as deletion of the mitochondrial pyruvate carrier (MPC1) in the neuroretina results in retinal degeneration^42^. Several findings have also implicated metabolic dysregulation in the pathogenesis of AMD^43–45^. In this disease, RPE dysfunction is considered a primary event, and studies have shown that RPE stability and its ability to support retinal function is reduced when it is forced to rely on glycolysis^46^. These data further support the hypothesis that metabolic imbalances, especially those in glucose metabolism, may underlie or enhance retinal degenerations.

The promotion of metabolic pathways in the retina has been shown to prevent retinal degeneration. Knockout of SIRT6 in *Pde6*-associated RP rods preserved retinal function and structure, which was associated with increases in expression of glycolytic proteins in the retina^8^. Furthermore, stimulation of the mTOR pathway with insulin treatment resulted in enhanced survival of cones, increased uptake and utilization of glucose, and elevated levels of NADPH in a fast-progressing model of RP^47^. Finally, our own studies have shown that activation of AMPK with metformin could preserve retinal function in both inherited and induced retinal degeneration models and was associated with increased expression of metabolism-related genes ^15^. These findings provide further evidence that enhancing metabolic activity, and in particular, glycolytic activity, may be protective in inherited retinal degenerations.

Studies have supported the idea that there is a distinct interdependence in the metabolism between RPE and photoreceptors. This has been corroborated in our present study. The RPE is known to utilize lactate to enhance glucose utilization in the retina. In the presence of reduced levels of lactate, RPE cells utilize more glucose for glycolysis^36^. Our metabolomic analysis revealed there was a significant reduction in lactate in the AMPK nDKO mice, which could reduce the amount that can be recycled back to the RPE for utilization. In addition to the loss of photoreceptor function, we observed RPE abnormalities as early as 4 months of age by TEM. Although AMPK was knocked out in the neuroretina and not the RPE, we hypothesize that the secondary loss of RPE function observed with ablation of AMPK in the neuroretina is due to decreased lactate production by the photoreceptors. This ultimately leads to RPE dysfunction, as the RPE are not able to maintain sufficient energy production. Although we hypothesized that AMPK may play an important role in cell-autonomous regulation of metabolism in the RPE, we did not observe any differences in retinal morphology or function with loss of AMPK in RPE cells. This may reflect the differences in metabolic utilization between the RPE and the photoreceptors. Further work is needed to investigate the role of AMPK in the RPE at older ages, and whether AMPK may be important to the RPE under stress conditions, when RPE may switch to a more glycolytic metabolism. Overall, this work establishes a significant role for AMPK in retinal metabolism and supports further investigation of AMPK and its energy-sensing pathways as therapeutic targets for maintaining metabolic homeostasis in retinal degenerations.

## Methods

RNA-sequencing data analysis: RNA-sequencing data was obtained from flow-sorted *Nrl-GFP* wild-type and *Pde6b*^*rd10*^ rods at age P19. Differential expression analysis was performed using edgeR on the raw counts of each biological sample^48^. Low-expressing genes were filtered out by Counts per Million (< 2 in all samples). Reference lists of metabolism and AMPK signaling genes were generated from the Gene Ontology (GO) resource^32,33^ and KEGG database^49–51^, respectively. A gene was considered differentially expressed if p < 0.05. A heatmap of differentially expressed AMPK genes was generated using the normalized counts with scaling by row using Heatmapper^52^. Clustering of genes and biological replicates was generated by Pearson Correlation.

Mice: 6 to 8-week-old BALB/cJ mice were purchased from The Jackson Laboratory and given 2 weeks to acclimate to the University of Florida animal facility. *Chx10-Cre* mice [Tg(Chx10-EGFP/cre,-ALPP)2Clc/J] were originally purchased from the Jackson Laboratory (stock # 005105) and crossed to the BALB/cJ strain for over 10 generations. Mice with floxed alleles for AMPKα1 (Prkaa1tm1.1Sjm/J) (stock # 014141) or AMPKα2 (Prkaa2tm1.1Sjm/J) (stock # 014142) were purchased from the Jackson Laboratory. These mice were backcrossed to BALB/cJ mice for at least 10 generations and mated to create AMPKα1ff;AMPKα2ff double knockout mice. These mice were bred to *Chx10-Cre* mice to create *Chx10-Cre*; AMPKα1ff;AMPKα2ff mice. Single knockouts were also bred to *Chx10-Cre* mice to generate either *Chx10-Cre*;AMPKα1ff mice or *Chx10-Cre*;AMPKα2ff mice. Mice were reared in 12-hour light 12-hour dark cyclic light with lights on at 6 AM. Light intensities in the cages were at 60 lux and food and water were given *ad libitum*. To rule out any potential confounding mutations, mice were screened for Rd1^53^, Rd8^54^, and Rpe65^55^ genes. All procedures were conducted in accordance with the Association for Research in Vision and Ophthalmology Statement for the Use of Animals in Ophthalmic and Vision Research and were approved by the University of Florida Institutional Animal Care and Use Committee.

Spectral-domain optical coherence tomography (SD-OCT): High resolution SD-OCT imaging was used to visualize retinal morphology (Bioptigen; Research Triangle Park, NC, USA). Mice were anesthetized with a solution of ketamine and xylazine. Eyes were dilated with drops of 1% tropicamide (AKORN Pharmaceuticals; Lake Forest, IL) and 2.5% phenylephrine (AKORN Pharmaceuticals; Lake Forest, IL) and to maintain corneal hydration artificial tears were used (Carboxymethylcellulose Sodium (0.5%), Glycerin (0.9%), Allergan; Dublin, Republic of Ireland). The Bioptigen Diver software (Bioptigen, Version 2.2) was used to manually measure outer nuclear layer (ONL) thickness using linear calipers to measure the ONL at pre-defined 125 um intervals.

Electroretinography (ERG): Mice were dark-adapted overnight and then were anesthetized and their eyes were dilated as described above. The Colordome ERG instrument (Diagnosys; Littleton, MA) was used to assess retinal function. Electrodes were inserted subcutaneously into the cheek (reference electrode) and in the tail (grounding electrode). Gold wire loops were placed on corneal surface with a drop of eye lubricant (Carboxymethylcellulose Sodium (0.5%), Glycerin (0.9%), Allergan; Dublin, Republic of Ireland. Stimulations were performed as described previously^9^.

Metabolomics: For steady-state metabolomics analysis, mice were subcutaneously injected with 300 mg/kg metformin (Sigma-Aldrich, PHR1084) for 7 days. Retinas were collected 2 hours after the last injection, rinsed with PBS, and flash-frozen in liquid nitrogen. Steady-state metabolites were measured by LC-MS as previously described^38,56,57^. For metabolic flux analysis, mice were IP-injected with 500 mg/kg ^13^C-labeled D-glucose (U-13C6, 99%, Cambridge Isotope Laboratories Inc.). Retinas were rapidly collected 45 minutes after injection using an established technique ^8^ in 200 µl cold HBSS, rinsed in PBS, and immediately flash frozen in liquid nitrogen. The retinas were then homogenized in methanol, chloroform, and water at a ratio of 700:200:50. The metabolites were dried, derivatized, and analyzed by GC-MS (Agilent 7890/5975C) and the chromatograms were analyzed using Agilent ChemStation software. The IsoCor software was used to correct the distribution of the mass isotopologues based on the natural abundance of isotopes. The abundance of the labeled ions over the total ion intensity was calculated.

qRT-PCR: For RNA: Retinas were collected into the Trizol reagent (Invitrogen, 15596026). According to the manufacturer’s protocol, the RNA was extracted and the iScript reverse transcription mix was used (Bio-Rad; Hercules, CA) to obtain cDNA templates. For Real-Time PCR: Primers were designed as described^9^. Ssofast SYBR Green Supermix (Bio-Rad; Hercules, CA) was used according to the manufacturer’s instructions and as described^9^. Expression was normalized to the housekeeping gene *Rpl19* and fold change was calculated relative to the control. Primers are listed in Supplementary Table 5.

Immunohistochemistry: Immunohistochemistry (IHC) was performed as previously described^9^. In brief, eyecups were collected, lightly fixed, and placed through a sucrose gradient. Frozen sections of 16 µm thickness were obtained.

Antibodies: Zonula occludens (ZO-1) (1:400 dilution, rabbit, polyclonal, Life Technologies). Electron Microscopy: Cervical dislocation was used to euthanize mice. After enucleation, eyes were dissected in 2.5% glutaraldehyde in 0.1 M phosphate buffer (pH 7.4) fixative. Tissue preparation and imaging were performed as described previously^58^.

Statistics: Statistical analysis was performed using GraphPad Prism (La Jolla, CA). To assess differences between two groups, paired or unpaired t-tests were used. To assess differences among more than 2 groups, analysis of variance (ANOVA) with a Tukey’s post hoc test was used. P-values less than 0.05 were considered statistically significant.

## Acknowledgments

The authors wish to acknowledge their funding sources, including the NIH R01 EY016459, the Foundation Fighting Blindness, and an unrestricted grant from Research to Prevent Blindness. The authors wish to acknowledge the contributions of Lena Phu for her assistance in genotyping, qRT-PCR, and measuring ONL thicknesses. The authors also wish to acknowledge the staff at the Robert P. Apkarian Integrated Electron Microscopy Core at Emory University for their assistance with the electron microscopy.

## Supplemental Figures

**Supplemental Figure 1.**
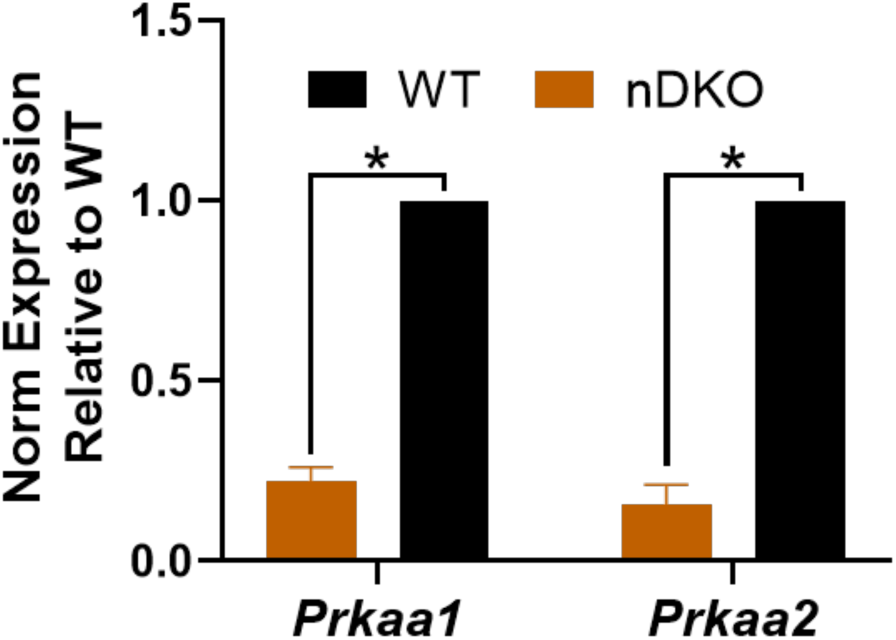
Deletion efficiency of AMPK nDKO mice. Expression of the genes encoding for the AMPK α1 and α2 subunits (*Prkaa1* and *Prkaa2*) in the neuroretina of nDKO mice relative to the WT. Note: This is consistent with efficient knockout in most retinal neurons, with residual expression coming from the non-targeted cells, vasculature, astrocytes, and microglia. Unpaired t-test, P<0.000001, n=6 per group.

**Supplemental Figure 2.**
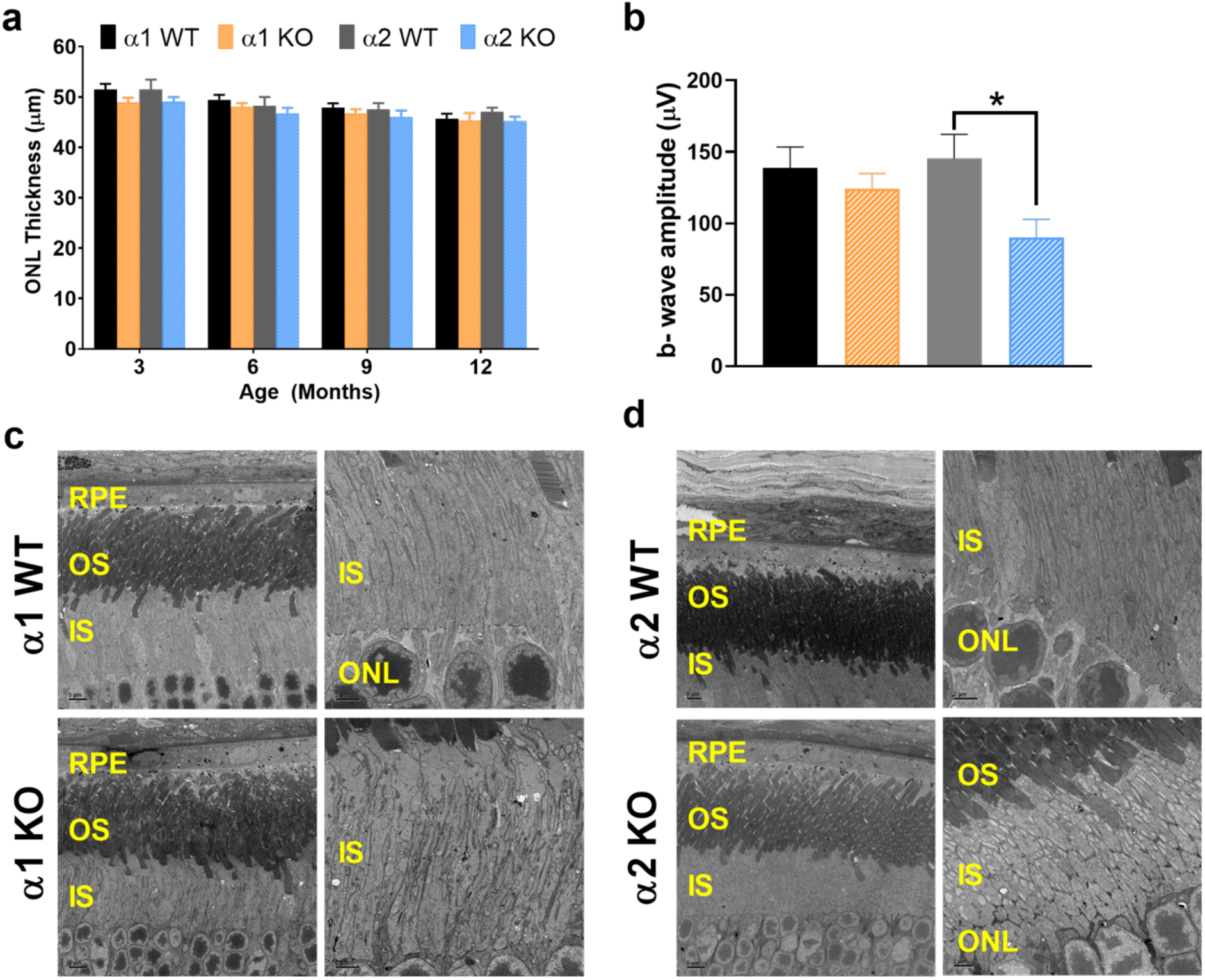
Knockout of single AMPK α1 or α2 subunit in the neuroretina does not affect retinal morphology, although the α2 subunit is required for cone photoreceptor function. a) ONL thicknesses from AMPK α1 single knockout mice compared to *Cre*-negative control WT mice over time from 2 months to 12 months of age. n=8-10 mice per group. b) Photopic ERG was used to measure cone photoreceptor function. ERG b-wave amplitudes were plotted comparing the single AMPK α1 or α2 knockouts to WT mice at 3 months of age. Loss of AMPK α2 activity led to a reduction in b-wave amplitude at light intensity 30 (cd.s/m^2^). n=6 mice per group. c-d) Mitochondria ultrastructure is maintained in the neuroretinas of single AMPK knockout mice. RPE, retinal pigmented epithelium; OS, outer segment; IS, inner segment.

**Supplemental Figure 3.**
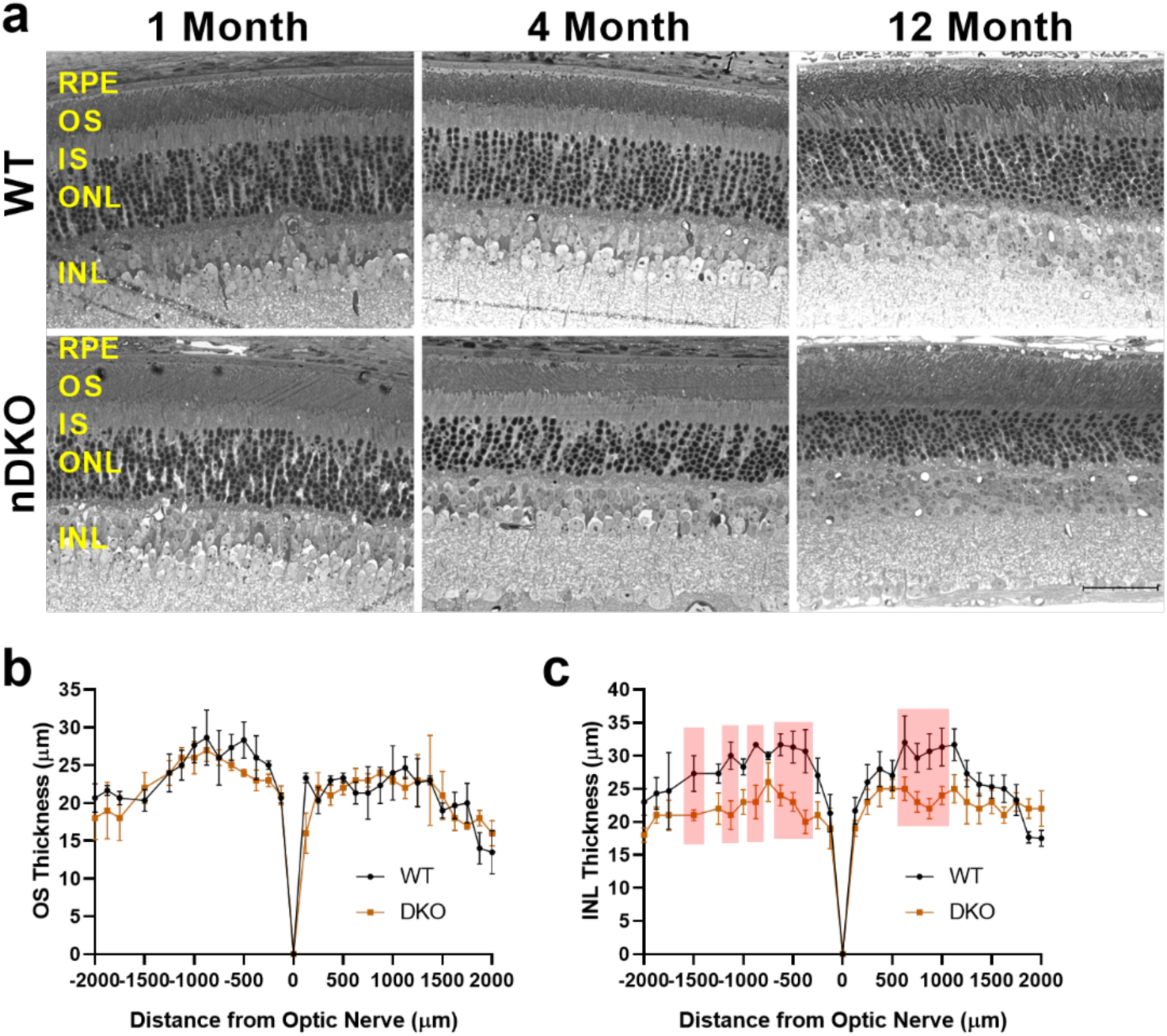
Alterations in retinal morphology with loss of AMPK. a) Representative histology images of retinal cross sections from WT (top) and AMPK nDKO (bottom) mice at 1 month (left), 4 months (middle), and 12 months (right) of age. Scale bar equals 50 µm. RPE = retinal pigment epithelium, OS = outer segments, IS = inner segments, ONL = outer nuclear layer, INL = inner nuclear layer. b-c) Quantification of OS (b) and INL (c) thicknesses from 12-month-old WT and nDKO retinas. Red shaded boxes indicate statistical significance. Multiple t-tests, n=3 WT and n=4 nDKO.

**Supplemental Figure 4.**
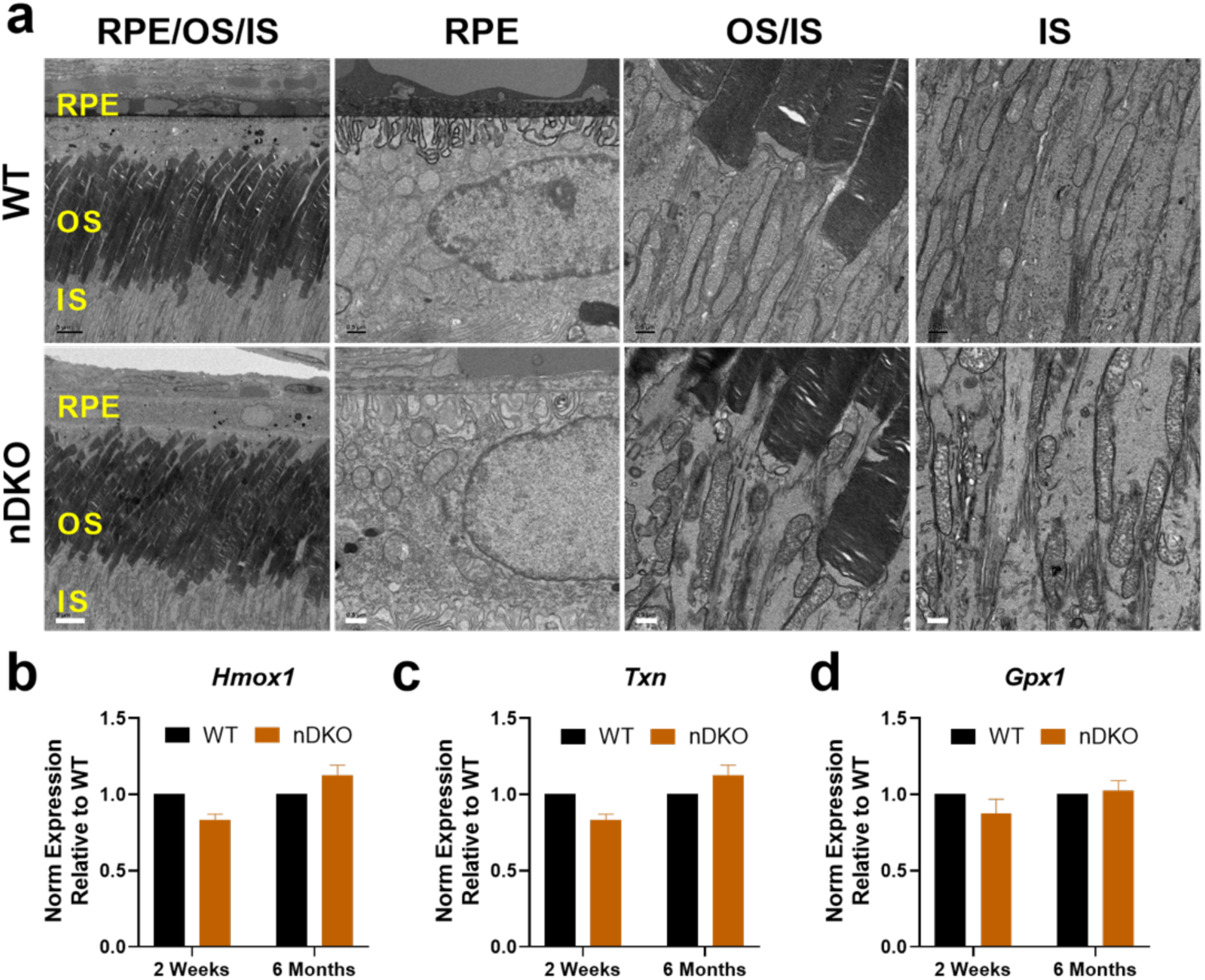
Retinal ultrastructure and oxidative stress-related gene expression. a) EM images from retinal cross sections of 1-month-old WT (top) and AMPK nDKO (bottom) mice. From left to right: RPE/OS/IS, RPE, OS/IS, and IS. Scale bars equal 5 and 0.5 µm. b-d) mRNA expression levels of the antioxidant genes *Hmox1* (b), *Txn* (c), and *Gpx1* (d) in the neuroretina of AMPK nDKO mice relative to WT at 2 weeks and 6 months of age. T-tests, P<0.05, n=3 per group.

**Supplemental Figure 5.**
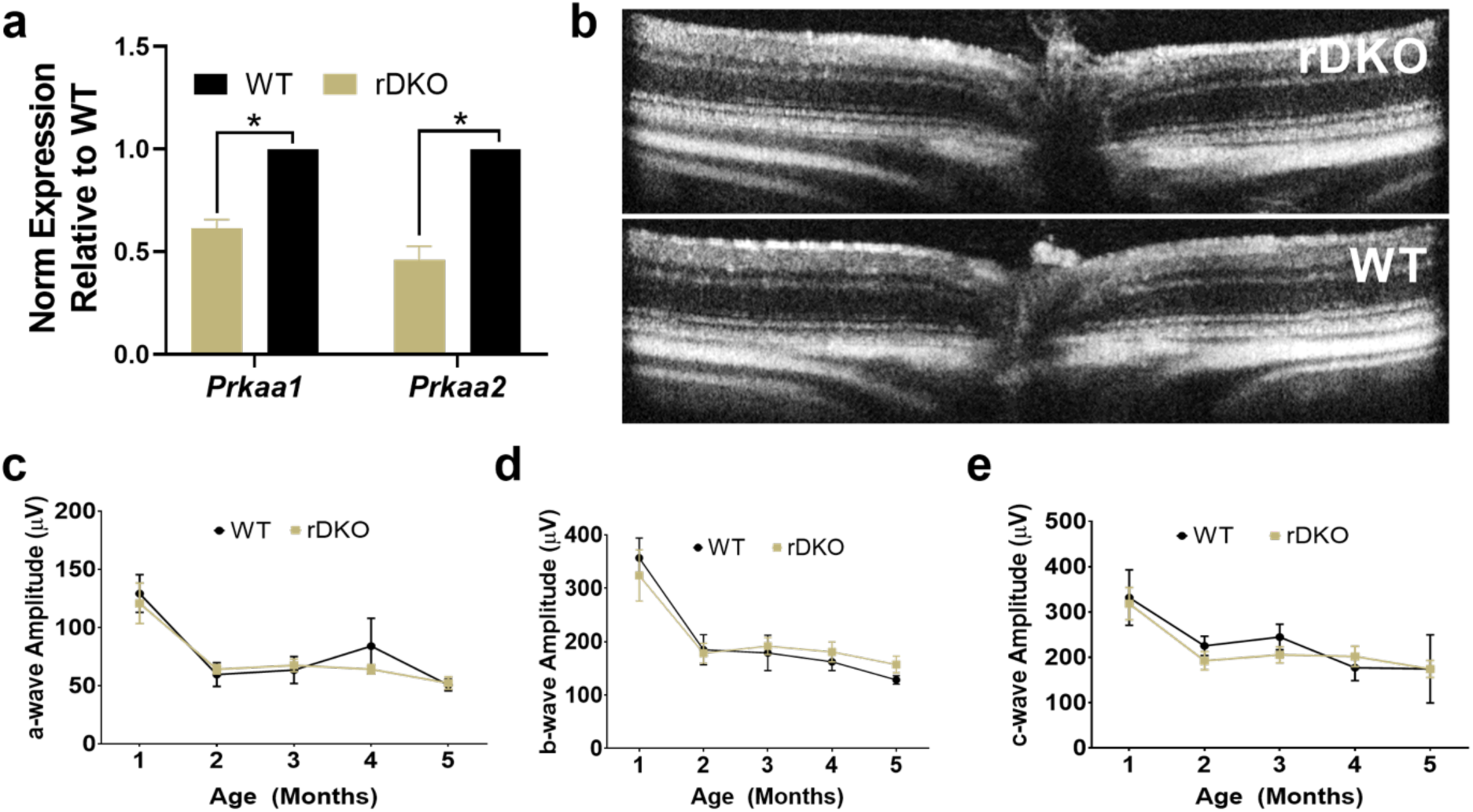
AMPK expression is not required for normal RPE function. a) Gene expression of the AMPK α subunits (*Prkaa1* and *Prkaa2*) relative to WT gene expression levels. Unpaired t-test, P<0.002, n=3 per group. Note: The deletion efficiency is likely higher than these values, but we collected the posterior eyecup, which contains a variety of cells including the RPE monolayer. Thus, there is likely contamination of other cells, such as extra-ocular muscle and the choroid that would increase levels of AMPK expression. Due to lack of specificity of commercially available antibodies for any of the AMPK subunits, we were not able to utilize other methods, such as IHC or western blotting, to confirm the deletion. b) Representative OCT images of 3-month-old AMPK rDKO (top) and WT (bottom) retinas showing no differences in morphology. c-e) Scotopic ERG a-wave corresponding to rod photoreceptor function (a), b-wave corresponding to inner retinal function (b), and c-wave corresponding to RPE function (c) with aging at 1.0 log (cd.s/m^2^) stimulation intensity. Two-way ANOVA, P>0.92, n≥4 per group.

